# Discerning Specific Thrombolytic Activities and Blood Clot Degradomes of Diverse Snake Venoms with Untargeted Peptidomics

**DOI:** 10.1101/2024.08.30.610527

**Authors:** Cara F. Smith, Mamadou Alpha Baldé, Lilyrose Bahrabadi, Merilyn Amponsah-Asamoah, Keira Y. Larson, Sean P. Maroney, David Ceja-Galindo, Martin Millimouno, Naby Camara, Jordan Benjamin, Nicklaus P. Brandehoff, Cassandra M. Modahl, Maxwell C. McCabe, Mitchell J. Cohen, Todd A. Castoe, Cellou Baldé, Kate Jackson, Stephen P. Mackessy, Kirk C. Hansen, Anthony J. Saviola

**Affiliations:** Department of Biochemistry and Molecular Genetics, 12801 East 17th Avenue, University of Colorado Denver, Aurora, CO, USA; Asclepius Snakebite Foundation, Aurora, CO, USA; Snakebite Center of Excellence, Institut de Recherche en Biologie Appliqueé de Guinée (IRBAG), Kindia, Guinea; Rocky Mountain Poison and Drug Center, Denver Health and Hospital Authority, Denver, CO, USA; Centre for Snakebite Research and Interventions, Liverpool School of Tropical Medicine, Pembroke Place, Liverpool, UK; Department of Surgery, University of Colorado Denver, 12631 East 17th Avenue, Aurora, CO, 80045-2527, USA; Department of Biology, The University of Texas Arlington, Texas, USA; Biology Department, Witman College, Walla Walla, WA, USA; Department of Biological Sciences, 501 20th Street, University of Northern Colorado, Greeley, CO 80639 USA

## Abstract

Identification and characterization of snake venom toxins that interfere with hemostasis have important implications for the treatment of snake envenomation, the bioprospecting of therapeutically useful molecules, and the development of research tools for investigating hematologic disorders. Many venoms have been shown to possess thrombolytic activity. However, it remains unclear if actions on other clot-stabilizing proteins beyond fibrin chains contribute significantly to venom-induced thrombolysis because the clot-wide targets of venom proteases and the mechanisms responsible for thrombolysis are not well understood. Here, we utilize a high-throughput time-based thrombolysis assay in combination with untargeted peptidomics to provide comprehensive insight into the effects of venom from six snake species on blood clot degradation. We compare thrombolytic profiles across venoms with variable levels of proteases and generate venom-specific fingerprints of cleavage specificity. We also compare the specific effects of venoms that possess a range of thrombolytic activity on fibrin subunits and other clot-bound proteins involved in clot structure. Venoms with higher thrombolytic activity demonstrated an enhanced ability to target multiple sites across fibrin chains critical to clot stability and structure, as well as clot-stabilizing proteins including fibronectin and vitronectin. Collectively, this study significantly expands our understanding of the thrombolytic and fibrinolytic effects of snake venom by determining the full suite of clot-specific venom targets that are involved in clot formation and stability.

## Introduction

Snake venom toxins that interfere with hemostatic processes have remained an enduring focus in snake venom research (1–4), and a myriad of procoagulant, anticoagulant, and fibrinolytic effects of various venoms and their toxins have been demonstrated experimentally and clinically across venomous snakes (5–7). Identification and characterization of these toxins and their activities have important implications for the treatment of snake envenomation, the bioprospecting of therapeutically useful molecules, and the development of research tools for investigating hematologic disorders (8). Though snake venoms are mixtures of both enzymatic and non-enzymatic toxins, that can act synergistically to produce a range of biological effects (9,10), much of the hematological disruption that results from snakebite is due to the action of enzymes belonging to the snake venom metalloprotease (SVMP) and snake venom serine protease (SVSP) families (11–15). These protease families are typically in high abundance in viper venoms and have a wide variety of target substrates in plasma, the extracellular matrix, and the basement membrane components of capillaries (13,14,16–20). Because of these activities, proteolytically rich venoms tend to promote defibrination, extravasation, capillary rupture, and hemotoxicity, resulting in local and/or systemic hemorrhage (12–14,21), and represent key contributors to human morbidity and mortality associated with snakebite worldwide (5,22).

In addition to pro- and anticoagulant properties, snake venoms and isolated venom proteins have demonstrated thrombolytic activity *in vitro* and *in vivo* (23–26), and many proteins involved specifically in thrombus formation and stability, including fibrinogen, fibronectin, and other structural proteins, coagulation factors, complement components, and plasminogen, have been identified as substrates of venom toxins (16–19,27). However, it remains unclear if the activity of snake venoms on other clot-stabilizing proteins beyond fibrin chains contributes significantly to the thrombolytic activity of snake venoms because the clot-wide targets of venom proteases and the mechanisms responsible for thrombolysis have yet to be fully investigated.

A detailed understanding of full protease substrate profiles (degradomes) is critical for an understanding of their overall function and possible contributions to disease pathophysiology (28). Recently, peptidomic approaches have gained traction in the study of disease progression and the discovery of disease state biomarkers (29). Peptidomics involves studying all peptides resulting from proteolysis and characterizing specific cleavage sites to understand enzyme-substrate target specificity. Peptidomics has been utilized in the discovery of novel substrates of previously characterized proteases (30) and the detection of disease biomarkers and predictors of clinical outcomes (31,32). Accordingly, peptidomic approaches can be utilized as a valuable complement to proteomics and facilitate the linkage of peptide-level changes with physiological responses, disease state, and pathophysiology (33).

The broad motivation of this study was to gain new insights into the dynamics of venom-induced thrombolysis through detailed characterization of the clot degradome produced by snake venom, and how this might vary across different species. We utilize a high-throughput time-based thrombolysis assay in combination with untargeted peptidomics to provide a comprehensive assessment of the effects of venom from six snake species on blood clot degradation. First, we compare thrombolytic profiles across venoms with variable levels of proteases and generate venom-specific fingerprints of cleavage specificity. We then identify and compare the specific effects of highly thrombolytic and less thrombolytic venoms on fibrin subunits and other clot-bound proteins involved in clot structure. Our findings provide critical new insight for understanding of the mechanisms responsible for thrombolysis and other hemostatic events triggered by snake venom proteases and demonstrate the power of protease substrate profiling for identifying the distinct activities and pathophysiological effects of venom toxins.

## Methods

### Halo thrombolytic assay

Blood samples were collected from healthy adult volunteers who gave informed consent to participate in the study. Human blood was collected into tri-sodium citrate (3.2%) tubes via venipuncture and clot formation was induced with the recombinant tissue factor Dade® Innovin®. Innovin® was reconstituted into 10 mL of ddH_2_O according to the manufacturer’s instructions. A clotting mixture of 15% (v/v) Innovin® and calcium chloride (67 mM) in HEPES buffer (25 mM HEPES, 137 mM NaCl) was prepared as previously described (34). Five μl of clotting mixture was pipetted onto the bottom edge of a flat-bottomed, tissue culture-treated 96-well plate. This clotting mixture was spread in a circular motion around the edge of the well before the addition of 25 μl of blood to form halos (Figure 1). A positive control that lacked the clotting mixture was included. The plate was incubated for 1 hour at 37°C to form stable clots.

**Figure 1.**
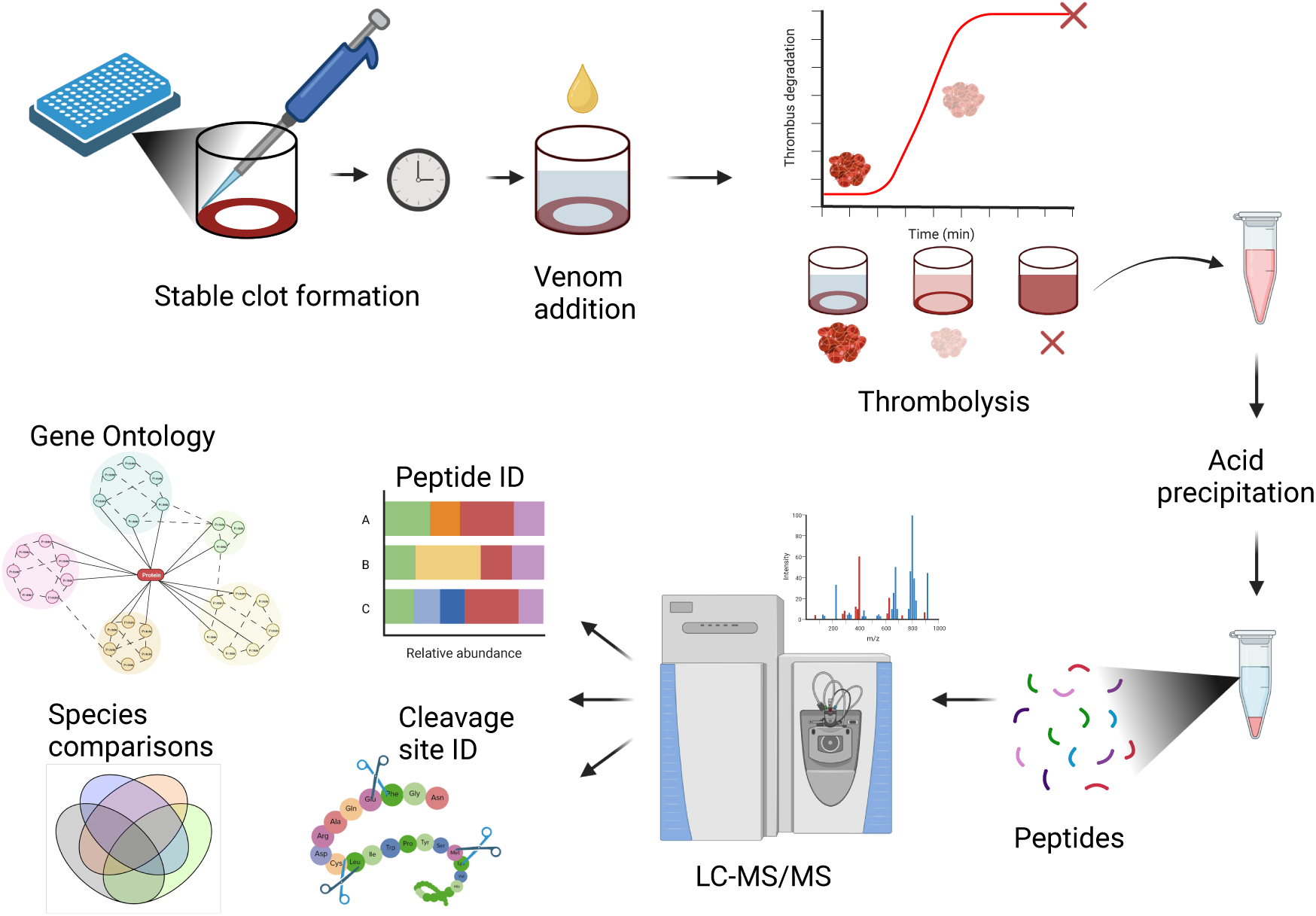
Halo thrombolytic assay and peptidomic analysis workflow. Whole blood clots are created in a 96-well plate. Snake venom or PBS was added after one hour and clot degradation was measured spectrophotometrically every minute for two hours. Supernatants from each well were removed and degradation peptides were purified with a PCA precipitation for LC-MS/MS analysis. Blood clot degradation products were identified and compared across species to determine major cleavage site patterns, protein targets, and target functions.

Fibrinolysis was tested with six different species of adult venomous snakes: *Bitis arietans* (BIAR), *Crotalus atrox* (southern Arizona; CRAT), *C. v. viridis* (northern Colorado; CRVI), *Dendroaspis viridis* (DEVI), *Naja savannula* (NASA), and *N. nigricollis* (NANI). Lyophilized venoms from CRAT and CRVI were obtained from snakes housed at the University of Northern Colorado (UNC) Animal Facility (Greeley, CO). All other venoms were obtained from the Research Institute of Applied Biology of Guinea (Kindia, Guinea) from snakes captured in the region. Venom was reconstituted at 4 mg/mL in ddH_2_O and diluted into 70 μl PBS for a final concentration of 1 mg/mL, 100 μg/mL or 10 μg/mL. Human plasmin diluted into PBS (0.01 U/mL, 0.3 U/mL, and 0.5 U/mL final concentration) was used as a positive control for fibrinolysis. Seventy μl of PBS + venom, PBS + plasmin, or PBS alone (negative control) was added in triplicate simultaneously into the wells containing the clots.

Clot degradation was measured every minute over 2 hours with a Tecan Spark® microplate reader by measuring absorbance change at 510 nm as previously described (Figure 1; (50)). Before each absorbance reading, the plate was orbitally shaken for 5 seconds at 180 rpm with a 3 mm amplitude. The percentage of degradation (Dx(t)) was calculated as previously described (34) by subtracting the negative control value and normalizing absorbances to the positive control. Positive controls (containing no clotting mixture) represented full degradation and negative controls (containing clotting mixture + PBS with no venom or plasmin) represented no fibrinolysis. The maximum clot lysis rate (CLR_max_) was determined by calculating the first derivative of degradation profiles in GraphPad Prism 10. The time elapsed until 50% lysis (T0.5) and the area under the curve (AUC) was also calculated based on the degradation curves generated in Prism.

### Venom comparisons

Fifty μg of venom was digested with trypsin for shotgun proteomics analysis as previously described (35). The volume corresponding to 50 μg was dried in a speed vacuum and resuspended in 8 M urea/0.1 M Tris (pH 8.5) and reduced with 5 mM TCEP (tris (2-carboxyethyl) phosphine) for 20 min at room temperature. Samples were then alkylated with 50 mM 2-chloroacetamide for 15 min in the dark at room temperature, diluted 4 times with 100 mM Tris-HCl (pH 8.5), and trypsin digested at an enzyme/substrate ratio of 1:20 overnight at 37°C. To stop the reaction, samples were acidified with 10% formic acid (FA), and digested peptides were purified with Pierce™ C18 Spin Tips (Thermo Scientific) according to the manufacturer’s protocol. Samples were dried in a speed vacuum and redissolved in 0.1% FA.

### Nano liquid chromatography-tandem mass spectrometry (LC-MS/MS)

LC-MS/MS was performed using an Easy nLC 1000 instrument coupled with a Q-Exactive™ HF Mass Spectrometer (both from ThermoFisher Scientific). Approximately 3 μg of digested peptides were loaded on a C_18_ column (100 μm inner diameter × 20 cm) packed in-house with 2.7 μm Cortecs C_18_ resin, and separated at a flow rate of 0.4 μL/min with solution A (0.1% FA) and solution B (0.1% FA in ACN) under the following conditions: isocratic at 4% B for 3 min, followed by 4%-32% B for 102 min, 32%-55% B for 5 min, 55%-95% B for 1 min and isocratic at 95% B for 9 min. Mass spectrometry was performed in data-dependent acquisition (DDA) mode. Full MS scans were obtained from m/z 300 to 1800 at a resolution of 60,000, an automatic gain control (AGC) target of 1 × 10^6^, and a maximum injection time (IT) of 50 ms. The top 15 most abundant ions with an intensity threshold of 9.1 × 10^3^ were selected for MS/MS acquisition at a 15,000 resolution, 1 × 10^5^ AGC, and a maximal IT of 110 ms. The isolation window was set to 2.0 m/z and ions were fragmented at a normalized collision energy of 30. Dynamic exclusion was set to 20 s.

### Analysis of mass spectrometry data

Fragmentation spectra were interpreted against an in-house protein sequence database generated from the venom gland transcriptome of each species using MSFragger within the FragPipe computational platform (36,37). Reverse decoys and contaminants were included in the search database. Cysteine carbamidomethylation was selected as a fixed modification, oxidation of methionine was selected as a variable modification, and precursor-ion mass tolerance and fragment-ion mass tolerance were set at 20 ppm and 0.4 Da, respectively. Fully tryptic peptides with a maximum of 2 missed tryptic cleavages were allowed and the protein-level false discovery rate (FDR) was set to < 1%. Identified proteins were classified by major venom toxin family and the relative abundance of major toxin families was compared across samples using sum-normalized total spectral intensity (38).

### Peptidomics

To capture the peptides released from lysed clots, we took the supernatant containing the products of clot degradation from each well after the 2-hour thrombolysis incubation and centrifuged for 15 minutes at 4°C and 18k x g. The soluble material was removed from the pellet and deproteinization with an acid precipitation was performed on the supernatant. Four molar ice-cold perchloric acid (PCA) was added to supernatants to a final concentration of 1M and vortexed. Samples were incubated on ice for 5 min and centrifuged at 18k x g for 2 minutes at 4°C. Supernatant was transferred to new tubes and neutralized with ice-cold 2M KOH, vortexed, and pH was checked with pH paper to ensure samples were between 6.5 – 8.0 pH. Neutralized samples were spun at 18k x g for 15 min at 4°C again and supernatant collected. Peptides were purified with Pierce^TM^ C_18_ Spin Tips (Thermo Scientific) according to the manufacturer’s protocol and LC-MS/MS was performed as described above using an Easy nLC 1000 instrument coupled with a Q-Exactive™ HF Mass Spectrometer.

Peptide spectra were interpreted against the UniProt human database (containing reverse decoys and contaminants) using MSFragger within FragPipe (36,37) as mentioned above. Protease specificity was set to non-specific and the FDR was set to < 1%. Principal component analysis and hierarchical clustering of identified proteins associated with each species and the Partial Least-Squares Discriminant Analysis (PLS-DA), variable importance in projection (VIP) plot, and volcano plot comparing thrombolytic to non-thrombolytic venoms were generated using MetaboAnalyst 5.0 (39). The volcano plot was generated with a fold change threshold of 3 (log_2_FC 1.58) and a p-value threshold of 0.05. Gene ontology analysis for the top 20 pathways associated with identified clot peptides was generated with ShinyGO 0.741 software using a 0.3 edge cutoff and a 0.05 FDR with the Biological Process database. Peptidomic data and protein-level cleavage patterns were visualized using Peptigram (40). PICS analysis of cleavage site and flanking amino acids specificity was performed with WebPICS (41). Protein networks were produced in STRING V12.0, where line weight indicates the strength of data support for protein-protein interactions and a minimum interaction score of 0.7. STRING protein network nodes were colored based on Reactome database categories.

## Results

### Halo assay fibrinolysis profiles

Thrombolysis profiles were generally dose-dependent in both venoms and plasmin (Figure 2a-g). At the high dose of plasmin, full degradation was reached by minute 42 (Figure 2g) but in BIAR and CRAT (Figure 2a-b) full degradation was not reached until minute 73 in all replicates. The highest dose of CRVI approached 50% clot lysis after 2 hours (Figure 2c), and all elapids showed minimal thrombolysis (Figure 2d-f). DEVI in particular appeared to have no detectable thrombolysis (Figure 2f), and AUC, T0.5, and CLR_max_ could not be calculated for this species, so it was omitted from further analyses.

**Figure 2.**
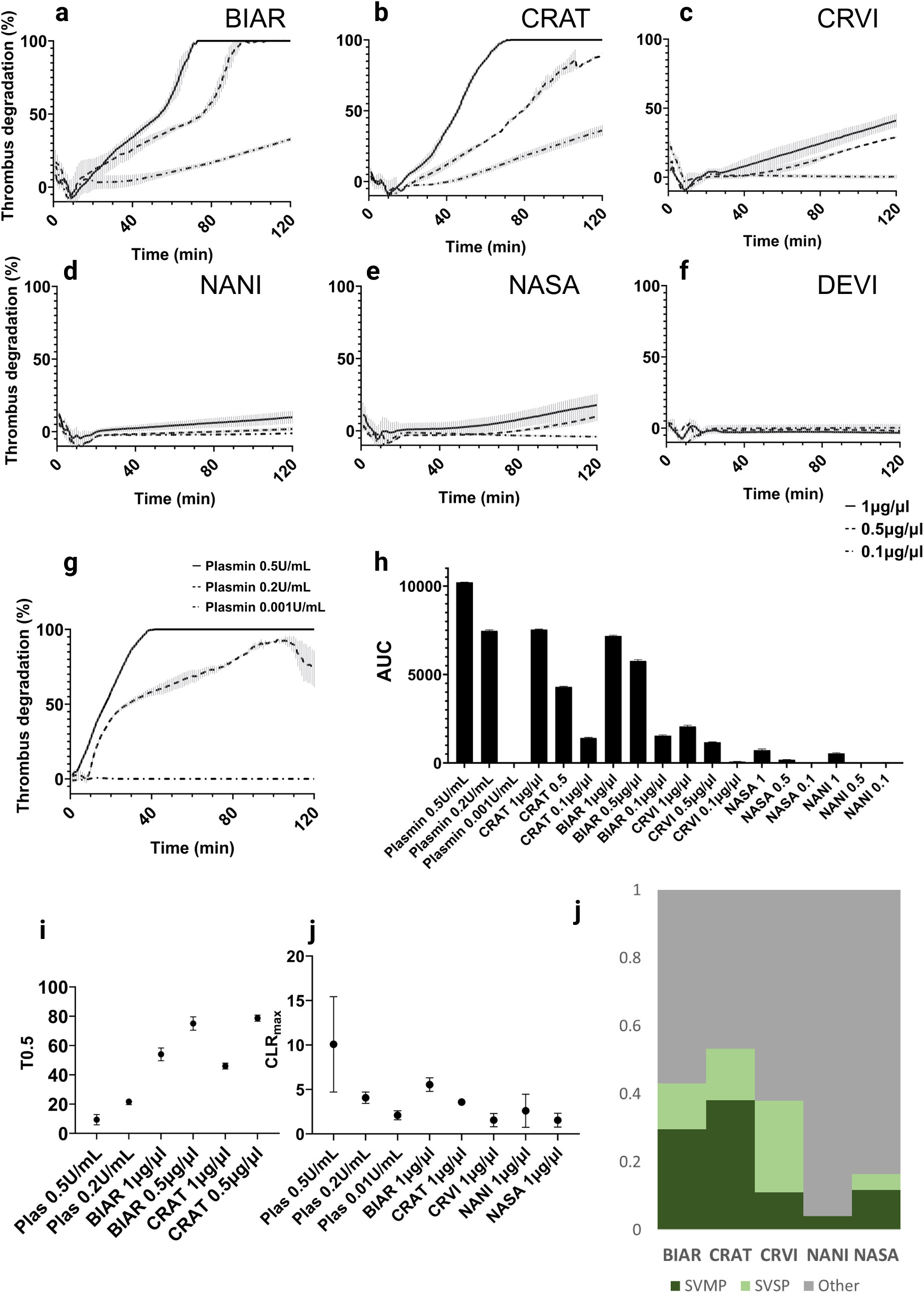
Snake venom thrombolytic profiles. Thrombolysis profiles of venom at 1μg/μl, 0.5μg/μl, and 0.1 μg/μl from a) *B. arietans* (BIAR), b) *C. atrox* (CRAT), c) *C. v. viridis* (CRVI), d) *N. nigricollis* (NANI), e) *N. savannula* (NASA), and f) *D. viridis* (DEVI). g) Thrombolysis profile of plasmin at 0.5U/mL, 0.2U/mL, and 0.001U/mL. h) Area under the curve (AUC) comparisons of thrombolytic profiles between species and across concentrations represented as mean ± SD i) Time to 50% lysis (T0.5) comparisons for all treatments that reached at least 50% lysis j) Maximal clot lysis rates comparing the highest concentration of venom (1μg/μl) used for all species to all three concentrations of plasmin. k) Shotgun proteomic data of venoms comparing relative abundance across species of the proteolytic enzymes snake venom metalloproteases (SVMPs) and snake venom serine proteases (SVSPs) to all other toxin families.

AUC also differed in a dose-dependent manner in plasmin and venoms with the highest AUC value found in 0.5U/mL plasmin (10,211 ± 12.2; Figure 2h) and lowest at 0.001U/mL plasmin (10.7 ± 2.6). In venom, the highest AUC values were found in CRAT and BIAR when comparing species at any concentration. AUC in NASA and NANI were negligible compared to viperid values. However, CRVI had significantly lower AUC than BIAR and CRAT.

The highest dose of plasmin (0.5U/mL) reached 50% lysis twice as fast as 0.2U (9. 3± 3.5 minutes vs. 21.7 ± 2.1 minutes, p<0.0063; Figure 2i). Plasmin reached 50% degradation (9.3 ± 3.5 minutes) significantly faster than even the most degradative snake venoms, BIAR and CRAT (54 ± 4.4 and 46 ± 2, respectively; p<0.0001). Both BIAR and CRAT reached 50% lysis at the highest dose (1µg/µl) significantly faster than the middle dose (0.5µg/µl; BIAR 54± 4.3 minutes vs. 75 ± 4.6 minutes, CRAT 46± 2 minutes vs. 78.7 ± 2.1 minutes; p<0.0001).

CLR_max_ differed in a dose-dependent manner in plasmin (p=0.044, p=0.005) except for the middle dose compared to the lowest dose (p=0.93; Figure 2j). The CLR_max_ value of the highest dose of plasmin was significantly higher than the highest dose of all venoms, except for BIAR (p=0.20). While CLR_max_ did not differ between the highest doses of any venoms, the highest value was found in the maximum dose of BIAR venom (5.5 ± 0.76 min^-1^), and the lowest value was from NASA (1.5 ± 0.77 min^-1^).

### Venom Composition

Venom toxins identified by LC-MS/MS were grouped into major toxin families, which were further categorized as one of two major venom proteases (SVSP, SVMP) or as other for all non-proteolytic enzymes and all nonenzymatic toxins. The highest proportions of proteases belonged to the vipers, with BIAR and CRAT having the highest abundances of SVMPs and CRVI with the highest SVSP abundance (Figure 2j). NANI and NASA have overall lower levels of venom proteases, though NASA SVMP abundance was similar to CRVI.

### Protein and peptide identification

The number of peptides identified from degraded clots ranged from 0 in clots incubated with DEVI venom to 2810 peptides from clots incubated with BIAR venom. The highest number of peptides identified came from clots incubated with the venom of the two vipers that produced the highest rates of thrombolysis in the halo assay (2810 in BIAR and 2293 from CRAT; Figure 3a) and the lowest number of peptides resulting from incubation with CRVI venom (1364 peptides). This is mirrored in the total number of identified proteins, with the highest number of proteins identified in BIAR (Figure 3b; 170 proteins) and CRAT (130 proteins) treated clots and the lowest in CRVI (75 proteins). On average, clots incubated with viper venoms (BIAR, CRAT, CRVI) produced more than twice the number of clot-based peptides (2156 and 1002 peptides, respectively; Figure 3c) and proteins (125 vs. 68 proteins; Figure 3d) than those incubated with elapid venoms (NASA, and NANI). However, these differences were not statistically significant (p=0.26 and p=0.15, respectively). Diversity values were calculated as the amount of cleavages/number of proteins. Interestingly, CRVI had the highest diversity value (Figure 3e; 36), while NANI had the lowest diversity value (26).

**Figure 3.**
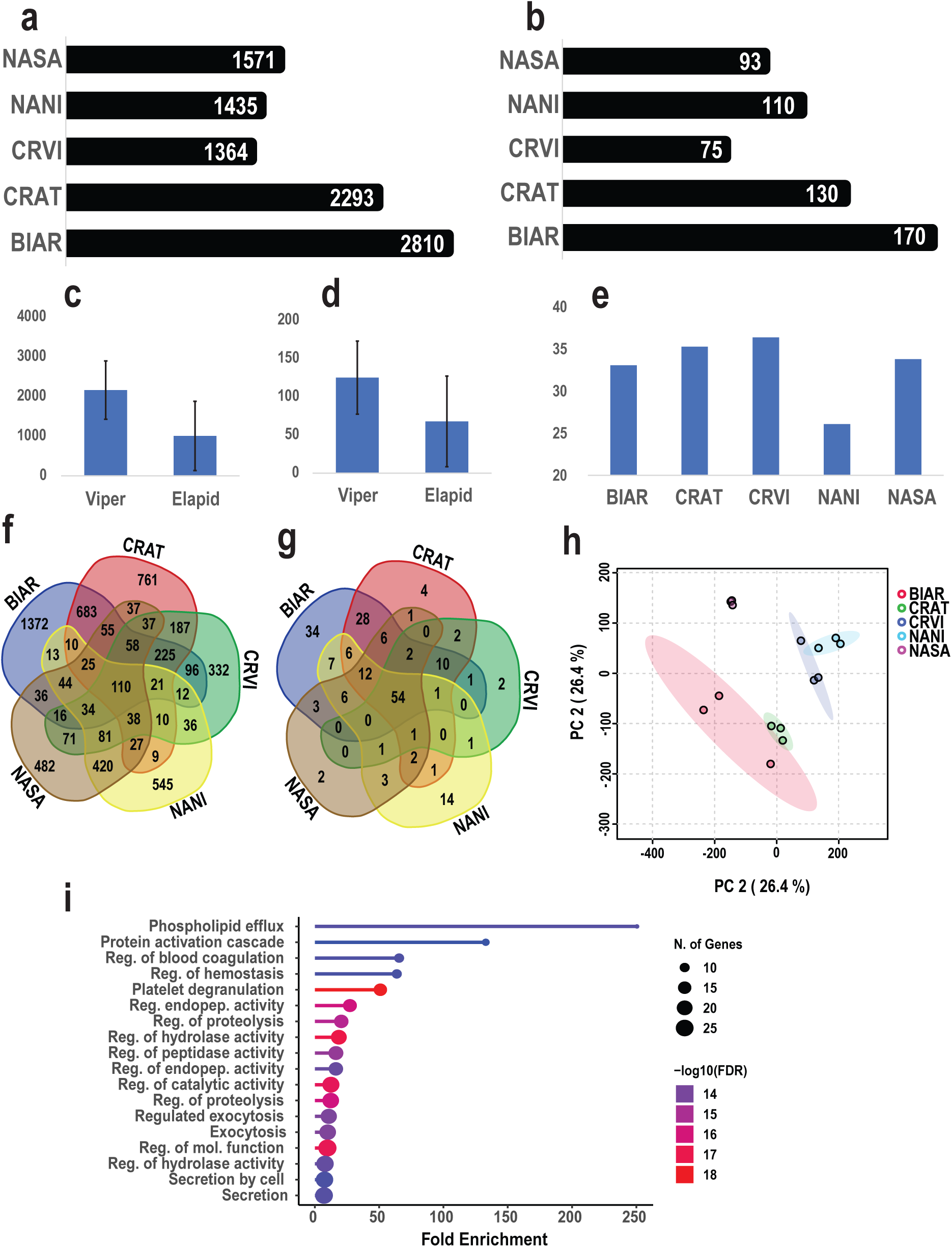
Peptide and protein identification of clot degradation products. The total number of unique a) peptides and b) proteins identified across species. Average number of c) peptides and d) proteins identified in vipers and elapids. e) Diversity values are calculated as the number of cleavages / the number of proteins identified. f) Venn diagram showing the number of unique and shared f) peptides and g) proteins across species. h) PCA plot with 95% confidence intervals of proteins identified in venom degradation profiles. i) Gene ontology analysis in ShinyGO 0.78 of the 54 proteins found in all degradation profiles.

There were 110 peptide degradation products mapping to 54 proteins common to all species (Figures 3f-g). Again, clots treated with BIAR venom produced the highest number of unique peptides and proteins (1372 unique peptides mapping to 34 proteins; Figures 3e-f), while the lowest number of unique peptides and proteins (332 and 2, respectively) came from clots incubated with CRVI venom.

Principle component analysis showed overall tight consistent clustering of replicates within species (Figure 3h). The two most thrombolytic venoms, BIAR, and CRAT-treated clot products clustered more closely together than with other species, while CRVI and NANI formed another distinct cluster. To identify common targeted pathways, we performed gene ontology (GO) analysis with the 54 proteins common to all clots (Figure 3g). The top 20 enriched pathways of common proteins were primarily involved in phospholipid efflux (n=7, Fold enrichment 251; Figure 3i), regulation of blood coagulation (n=10, Fold enrichment=66), and platelet degranulation (n=15, Fold enrichment=51).

### Fibrinogen cleavage profiles

All species showed more cleavage products of fibrin chain alpha than beta or gamma, and CRAT and BIAR showed the highest coverage/number of peptides detected overall (Figure 4a). All species demonstrated degradation products associated with the cleavage of fibrinopeptide A (residues 20-35) and regions between residues 500-600 spanning the αC domain of the main fibrinogen alpha chain (Figure 4b; Supplemental Figure 1). CRAT and BIAR also demonstrated high coverage and depth of chain alpha peptides in the αC Connector region between residues 200 to 400. The highest intensities came from the peptides ^223^LIKMKPVPDLVPGN^236^ and ^393^FRPDSPGSGNARPNNPDWGT^412^ in CRAT and ^225^KPVPDLVPGN^236^ in BIAR (Supplemental Figure 1). We identified the thrombin cleavage site at residues 35-36 across species and found degradation peptides flanking plasmin cleavage sites at residues 100-101 and 123-124 in BIAR and CRAT only. We also identified peptides flanking cross-linking sites at residues 322, 347, 385, 527, 558, 575, 581, and 599 in all species.

**Figure 4.**
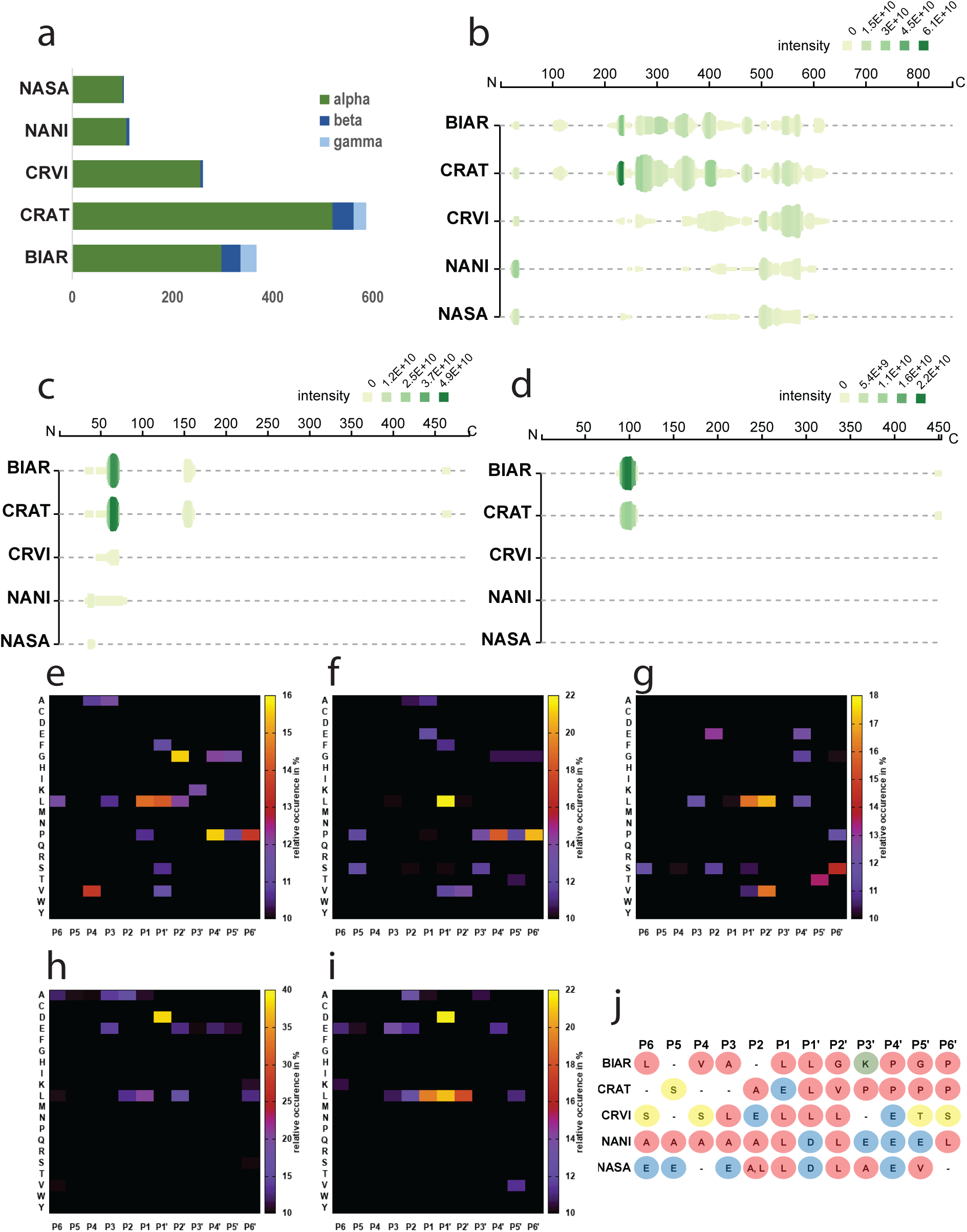
Peptidomic analysis of fibrin and cleavage specificity. a) Number of unique peptides belonging to the alpha, beta, or gamma chains of fibrin. Peptide cleavage patterns and depth of fibrin chain b) alpha, c) beta, and d) gamma represented with Peptigram (Manguy et al., 2017). The plot height represents the number of overlapping peptides at a position and the color represents sum of the intensities of the overlapping peptides. Heatmap of the relative occurrence (in %) of each amino acid residue among the top 400 most abundant peptides identified from clots treated with e) BIAR, f) CRAT, g) CRVI, h) NANI, and i) NASA venoms. j) Most preferred residue at each position from P6-P6’ for each species.

Relative to alpha, fibrinogen beta cleavage was significantly lower in all species but was particularly low in CRVI, NANI, and NASA (Figure 5c). BIAR and CRAT showed the areas of highest peptide coverage between 58-72 of the main beta chain, and another area between 150 and 163 that was absent in all other species, where two plasmin cleavage sites are found (Supplemental Figure 2). Venom of all species except CRVI liberated fibrinopeptide B (residues 31-44) and all species except NASA showed cleavage at the thrombin cleavage site (44–45). The peptide ^58^SLRPAPPPISGGGYR^72^ had the highest intensity in BIAR and CRAT, which was located downstream from a beta chain polymerization site at residues 45-47. Only BIAR and CRAT venoms produced degradation products of fibrinogen gamma from residues 66-109, which flank plasmin cleavage sites at 84-85 and 88-89 and from residues 446 to the C-terminus (Figure 4d; Supplemental Figure 3). The C-terminal peptides identified in BIAR and CRAT are downstream of a fibrin alpha-gamma binding site and gamma polymerization site at residues 400-422 and the platelet aggregation site at residues 423-437.

**Figure 5.**
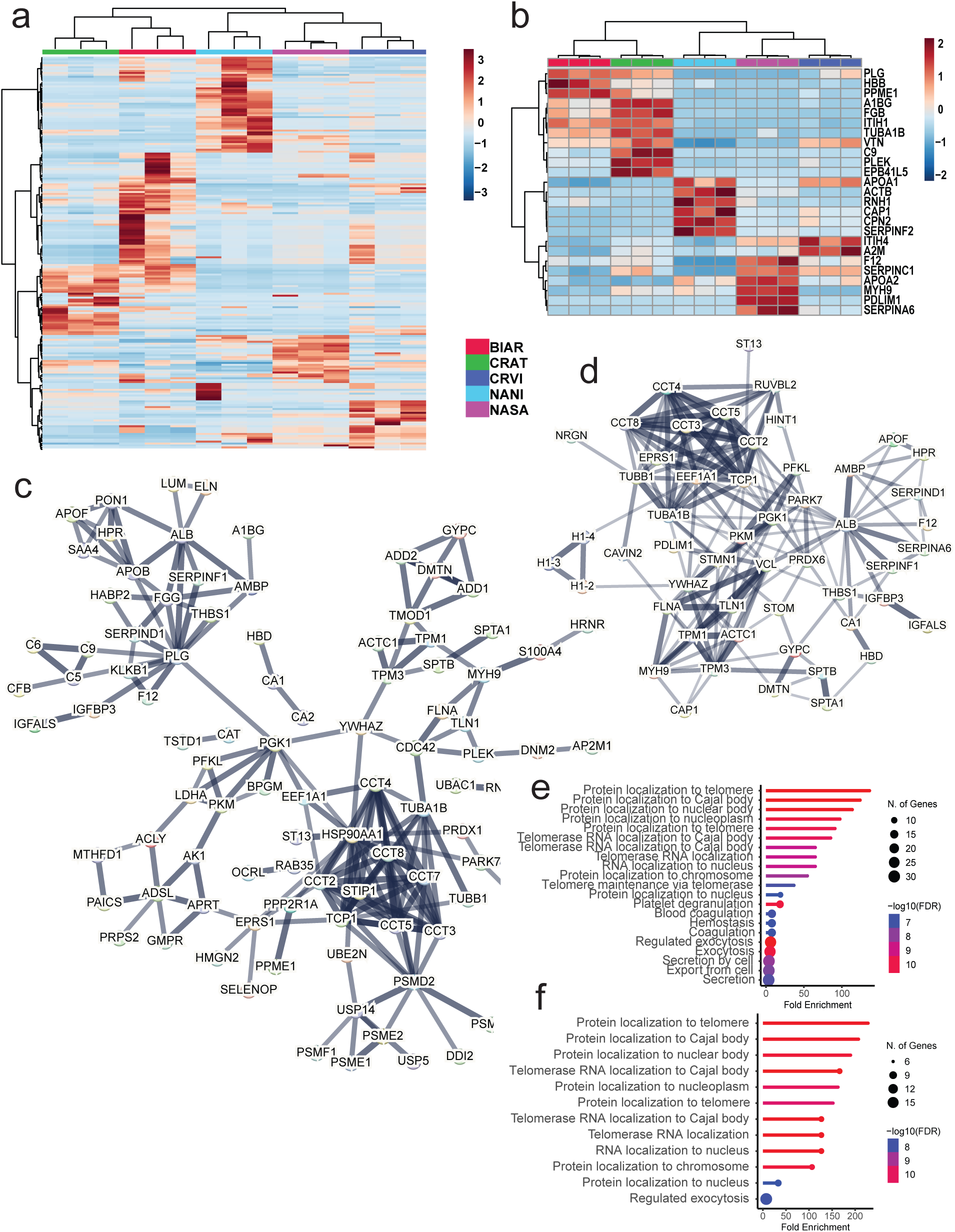
Species and family level protein identification and gene ontology comparisons. Heatmap of hierarchical clustering analysis of intensities from a) all proteins and b) top 25 proteins by p-value showing the clustering of triplicates within a species and of CRAT with BIAR and CRVI with NASA and NANI. Protein interaction networks generated in STRING V12.0 for clot degradation products shared among c) viper venoms and d) elapid venoms. Protein network line weight indicates the strength of data support for protein-protein interactions, at a minimum interaction score of 0.7. Gene ontology analysis in ShinyGO 0.78 of proteins identified in clot degradation products from e) vipers and f) elapids.

### Venom-wide cleavage sites

Fingerprints of cleavage specificity of the top 400 peptides by intensity revealed differences in both snake family- and species-level cleavage residue preferences. Leucine at P1’ was favored in viper degradomes (Figures 4e-g and 4j), particularly in CRAT, but there was also a preference for leucine at P2’ for CRVI and P1 in BIAR. Viperid venoms also showed a preference for valine at P2’ (CRVI; Figure 4g) and P4 (BIAR; Figure 4e). BIAR and CRAT showed a preference for proline in the P4’-P6’ region (Figures 4e-f). Last, BIAR showed a strong preference for glycine at P2’ (Figure 4e) and CRVI showed a preference for serine at P6’ (Figure 4g). NASA and NANI-induced cleavage events demonstrated a preference for aspartic acid at P1’ and a preference for leucine from P1-P2’ of NASA. (Figure 4h-j).

We determined overall consensus preferences at each site from P6-P6’ for each venom (Figure 4j). In general, we noted a strong preference for hydrophobic residues across flanking sites of all species, with a particular preference for leucine at P1 in all species but CRAT. All vipers preferred leucine at P1’ and both elapids preferred the residue aspartic acid at P1’. We also noted the higher preference for acidic residues aspartic acid and glutamic acid in both elapid species, and a preference for polar neutral residues in flanking regions in CRVI.

### Gene Ontology

Hierarchical clustering of all degradation products with sum normalized intensity shows that the two vipers BIAR and CRAT cluster together and are most similar, while CRVI clusters with the two elapids, NASA and NANI (Figure 5a). These comparisons also reveal different clusters of proteins that dominate the clot degradome of each species. When the top 25 proteins as determined by p-value are examined, the species-level cluster patterns remain consistent and the BIAR and CRAT degradome profiles closely resemble each other (Figure 5b). There also appear to be distinct suites of proteins that are uniquely higher in relative intensity across species. For example, the two most thrombolytic species, CRAT and BIAR, had higher relative intensities of plasminogen (PLG), hemoglobin (HBB), fibrinogen beta (FGB), vitronectin (VTN), and complement C9 (C9). NANI had a higher relative intensity of apolipoprotein A-I (APOA1), cytoplasmic actin (ACTB) and alpha-2-antiplasmin (SERPINF2), while NASA had a higher intensity of coagulation factor XIIa (F12), antithrombin-III (SERPINC1), and apolipoprotein A-II (APOA2). CRVI had a higher relative abundance of alpha-2-macroglobulin (A2M).

To identify snake family-level patterns in clot targets we mapped protein-protein interactions in STRING V12.0 for all degradation products identified in clots incubated with the viper venoms (CRVI, CRAT, and BIAR) and with the elapid venoms (NASA and NANI). The viperid protein network had a significant protein-protein interaction (PPI) enrichment p-value (p<0.0001) an average local clustering coefficient of 0.508 and an average node degree of 2.9 (Figure 5c). The proteins with the highest degree of connectivity (edges) were heat shock protein HSP 90-alpha (HSP90AA1; n=17), plasminogen (PLG; n=13), T-complex protein 1 subunit beta (CCT2; n=12), and T-complex protein 1 subunit alpha (TCP1; n=12). The elapid protein network had a significant PPI enrichment p-value (p<0.0001) an average local clustering coefficient of 0.42 and an average node degree of 5.56 (Figure 5d). The proteins with the highest degree of connectivity (edges) were albumin (ALB; n=17), phosphoglycerate kinase 1 (PGK1; n=16), and T-complex protein 1 subunit alpha (TCP1; n=15).

Based on analyses in ShinyGO of degradation products, the top 20 enriched pathways of viper-specific proteins were primarily involved in protein localization to telomeres (n=10, Fold enrichment 135), telomerase RNA localization (n=7, Fold enrichment=64), platelet degranulation (n=13, Fold enrichment=18), blood coagulation (n=15, Fold enrichment=8), and hemostasis (n=15, Fold change=8; Figure 5e). The top 20 enriched pathways in ShinyGO 0.741 of elapid-specific proteins were primarily involved in protein localization to telomeres (n=6, Fold enrichment 224), telomerase RNA localization (n=7, Fold enrichment=125), and protein localization to the nuclear body (n=6, Fold enrichment=187; Figure 5f).

### Differences between venoms with varying levels of thrombolytic activity

To identify the major defining characteristics between strongly thrombolytic and minorly thrombolytic venoms we created a volcano plot and performed PLS-DA to identify proteins with degradation products that significantly differed in abundance that might define the two groups. The volcano plot showed BIAR and CRAT clot degradomes produced significantly higher abundance of 74 proteins including tubulin alpha 1B chain (TUBA1B), fibrinogen beta and gamma chains (FGB and FGG), N-acetylmuramoyl-L-alanine amidase (PGLYRP2), vitronectin (VTN), ECM protein 1 (ECM1), inter-alpha-trypsin inhibitor heavy chain H1 (ITIH1), hemoglobin subunit delta (HBD), pleckstrin (PLEK), and fibronectin as major outliers (Figure 6a; Supplemental table 1). The PLS-DA VIP plot demonstrated that the top 3 components with the highest VIP scores included alpha-1-antitrypsin (SERPINA1), hemoglobin subunit beta (HBB), and fibrinogen alpha (FGA; Figure 6b). The top 15 VIP scores also included fibrinogen beta chain (FGB), alpha2-HS-glycoprotein (AHSG), glyceraldehyde-3-phosphate dehydrogenase (GAPDH), vitronectin (VTN), fibrinogen gamma (FGG), inter-alpha-trypsin inhibitor (ITIH1) and alpha-synuclein (SNCA; Figure 6b; Supplemental Table 2)

**Figure 6.**
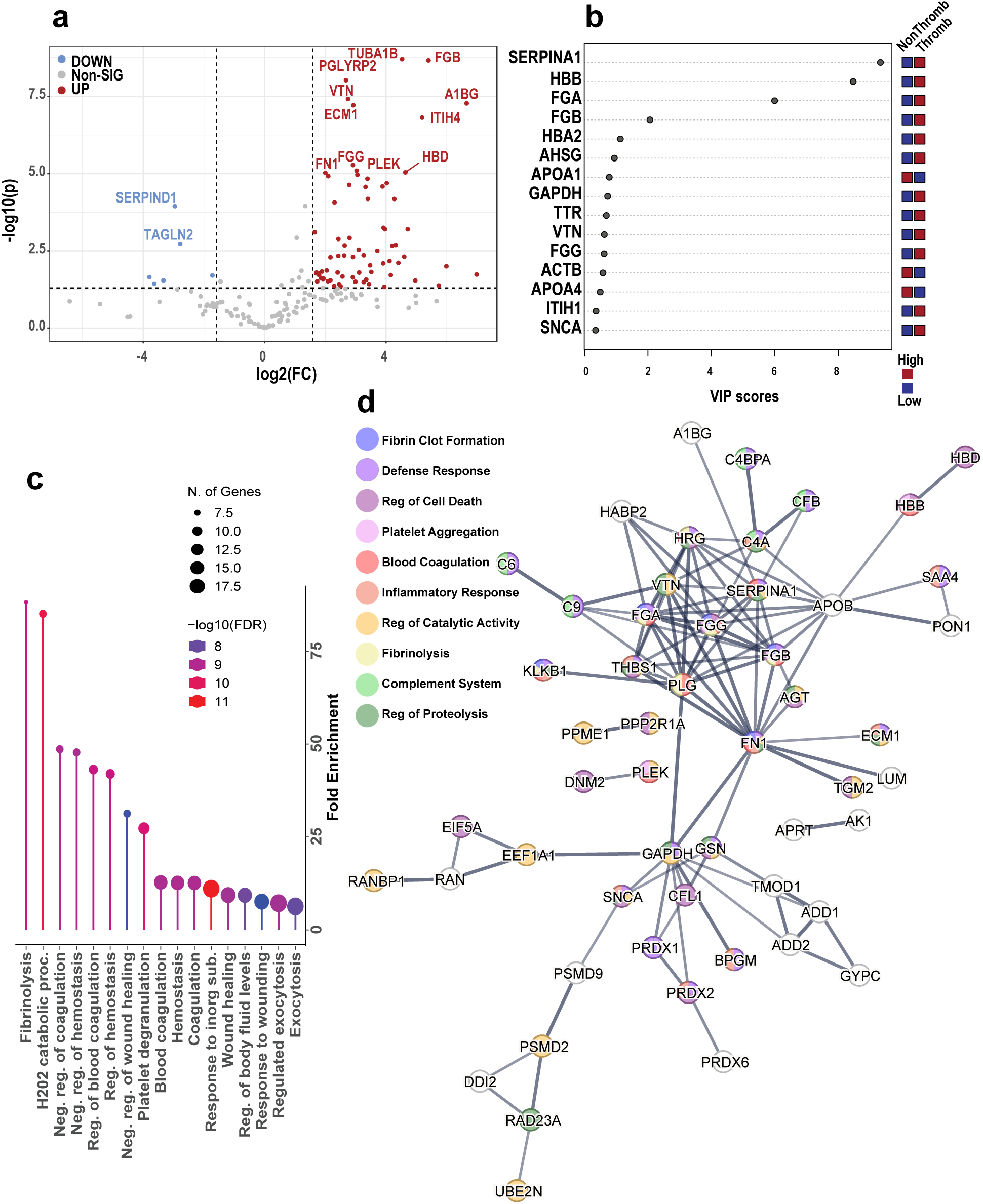
Defining features of thrombolytic venoms. a) Volcano plot comparing differences in protein abundance between clots incubated with thrombolytic venoms that reached 100% thrombus degradation (BIAR and CRAT) and less thrombolytic venoms that did not reach 50% degradation after two hours. b) VIP scores of the 15 most important proteins in PLS-DA analysis distinguishing thrombolytic from less thrombolytic venoms. Red indicates the group that had a significantly higher abundance of a protein and blue indicates the group that had a significantly lower abundance of the specified protein. c) Gene ontology analysis in ShinyGO 0.78 of proteins identified as significantly higher in thrombolytic venoms. d) STRING V12.0 protein interaction network with gene ontology analysis overlayed of proteins identified as significantly higher in thrombolytic venoms.

Shiny GO analysis of enriched clot degradome products produced by thrombolytic venoms identified in the volcano plot showed terms that are primarily related to hemostasis, including fibrinolysis (n=7; fold enrichment 88; Figure 6c), negative regulation of blood coagulation (n=8; fold enrichment = 48), and platelet degranulation (n=11; fold enrichment=27). Other pathways include hydrogen peroxide catabolic processes (n=8; fold enrichment=85), response to inorganic substance (n=19; fold enrichment=11), and regulation of body fluid levels (n=15; fold enrichment=9).

A STRING V12.0 PPI network of the degradation targets that were significantly higher in thrombolytic venoms produced a network with 71 nodes and 103 edges with an average node degree of 2.9, a PPI enrichment p-value <1.0e-16 and an average local clustering coefficient of 0.505 (Figure 6d). Major gene ontology biological processes identified included defense response (FDR=2.37e-06), regulation of catalytic activity (FDR=0.0084), regulation of cell death (FDR=0.0032), regulation of proteolysis (FDR=2.99e-05), inflammatory response (FDR=0.00092), blood coagulation (FDR=2.01e-06), fibrinolysis (FDR=9.1e-07), platelet aggregation (FDR=1.46e-05), fibrin clot formation (FDR=2.99e-05), and complement activation (FDR=0.00089). The nodes with the highest degree of connectivity were fibronectin (FN1; n=14), plasminogen (PLG; n=13), apolipoprotein B-100 (APOB; n=11), fibrinogen alpha chain (FGA; n=11), and fibrinogen beta chain (FGB; n=11).

## Discussion

Here, we demonstrate the utility of a high-throughput halo assay in the quantification of venom-induced clot degradation and compare the thrombolytic activities across five snakes with varied venom composition. Differences in clot lysis rates are associated with variable effects and cleavage patterns towards all fibrin chains and the clot-stabilizing proteins factor XIIIa, fibronectin, and vitronectin. Both the high abundance of proteolytic proteins and the ability to target multiple cross-linking, cell adhesion, and plasmin cleavage sites of fibrin, as well as structurally pivotal clot-bound proteins, are likely responsible for the high thrombolytic activity of BIAR and CRAT. Ultimately, this study expands our understanding of the thrombolytic and fibrinolytic effects of snake venom by determining the full suite of clot-specific venom targets that are involved in clot formation and stability. Our data adds to the repertoire of snake venom hematological targets and provides a holistic picture of the molecular basis of thrombolysis catalyzed by snake venoms, paving the way for further investigation of thrombolytic patterns of venom toxins using -omics technologies.

Vipers in general are well known to produce highly proteolytic venom compared to elapids (43,44). In this study, the three vipers demonstrated much higher overall protease abundance than the elapids; however, only the two highly thrombolytic viper species, BIAR and CRAT, produce venoms with higher levels of SVMPs, mirrored by the larger number of degraded peptides in their clot degradomes. Even the less thrombolytic venoms did produce proteolytic activity against fibrinogen alpha and beta chains and other clot-stabilizing proteins, suggesting that the higher rates of thrombolysis in CRAT and BIAR are, unsurprisingly and at least in part, dependent on the higher abundance of metalloproteases in their venoms.

Only two species, BIAR and CRAT, were able to produce full thrombolysis of clots after two hours. The third viper, CRVI, produced moderate thrombolysis at the highest concentration, but it failed to reach 50% clot lysis (T0.5) after two hours. Of the three elapids tested, only NANI and NASA produced slight thrombus degradation at the highest concentrations tested. DEVI failed to produce any detectable thrombolysis in the halo assay, mirrored by the lack of degradation peptides detected in the halo clot supernatants. This lack of observed clot degradation from DEVI is consistent with past studies noting that some *Dendroaspis* venoms possess anticoagulant properties but also inhibit clot lysis (42).

The generation of venom-induced cleavage fingerprints provides key new perspectives on the combined cleavage patterns induced by venom toxins, which has the potential to reveal both novel patterns of proteolysis and strengthen the ability to predict other unexplored substrates. Species showed characteristic differences in cleavage site preferences indicative of an overall divergence in activities between active proteases in each venom. As expected, viper venoms appeared to have an overall larger fingerprint than elapids, likely due to the overall higher abundance and complexity of venom proteases. However, despite using crude venom, proteolytic signatures resulting from the action of highly abundant SVMPs in viperid venoms were still detectable. For example, we detected the previously noted preference for Leu at P1’ of SVMPs (and metalloproteases in general) in all three viper venoms (16–18,45,46).

The detection of venom-specific cleavage events may be complicated by the indirect effects of envenomation via the activation of endogenous proteases by venom toxins (47,48). For example, elapids NASA and NANI demonstrated the meprin characteristic preference for Asp at the P1’ position, suggesting that the activation of endogenous proteases like meprin is responsible for some cleavage patterns detected rather than the direct action of venom proteases (49–51). These indirect cleavage events should be further elucidated by a thorough understanding of both venom and endogenous protease cleavage specificity across species and toxin protease families.

Previous *in vivo* studies have investigated the proteomic and peptidomic effects of envenomation using both biofluids and tissues. We identified many of the same proteins in the clot degradome that were previously identified either as venom protease targets or as effector proteins, including coagulation cascade factors, serpins, apolipoproteins, complement factors, ECM proteins, and fibrinogen chains (16–18,51,52) (Figure 7). Taken together, changes in the abundance of these endogenous proteins affect a wide variety of physiological responses to envenomation including thromboinflammation, immune system activation, hemostatic alterations, and cell and tissue destruction (16–19,51–53). However, correlating the differences in protein abundance of envenomated tissues to measurable differences in biological and physiological functions presents a significant challenge. As demonstrated here and in other studies, a number of the proteins identified as venom toxin targets, including coagulation factor XIIIa, fibronectin, vitronectin, alpha-2-antiplasmin, plasmin(ogen), thrombin, and alpha-2-macroglobulin comprise the main scaffold of clots or have direct effects on clot stability and/or thrombolysis. To understand how some of these venom targets relate specifically to venom-induced clot degradation, we focused specifically on integrating the observed differences in clot degradome peptides and proteins with the measurable differences in thrombolytic activity between snake venoms.

**Figure 7.**
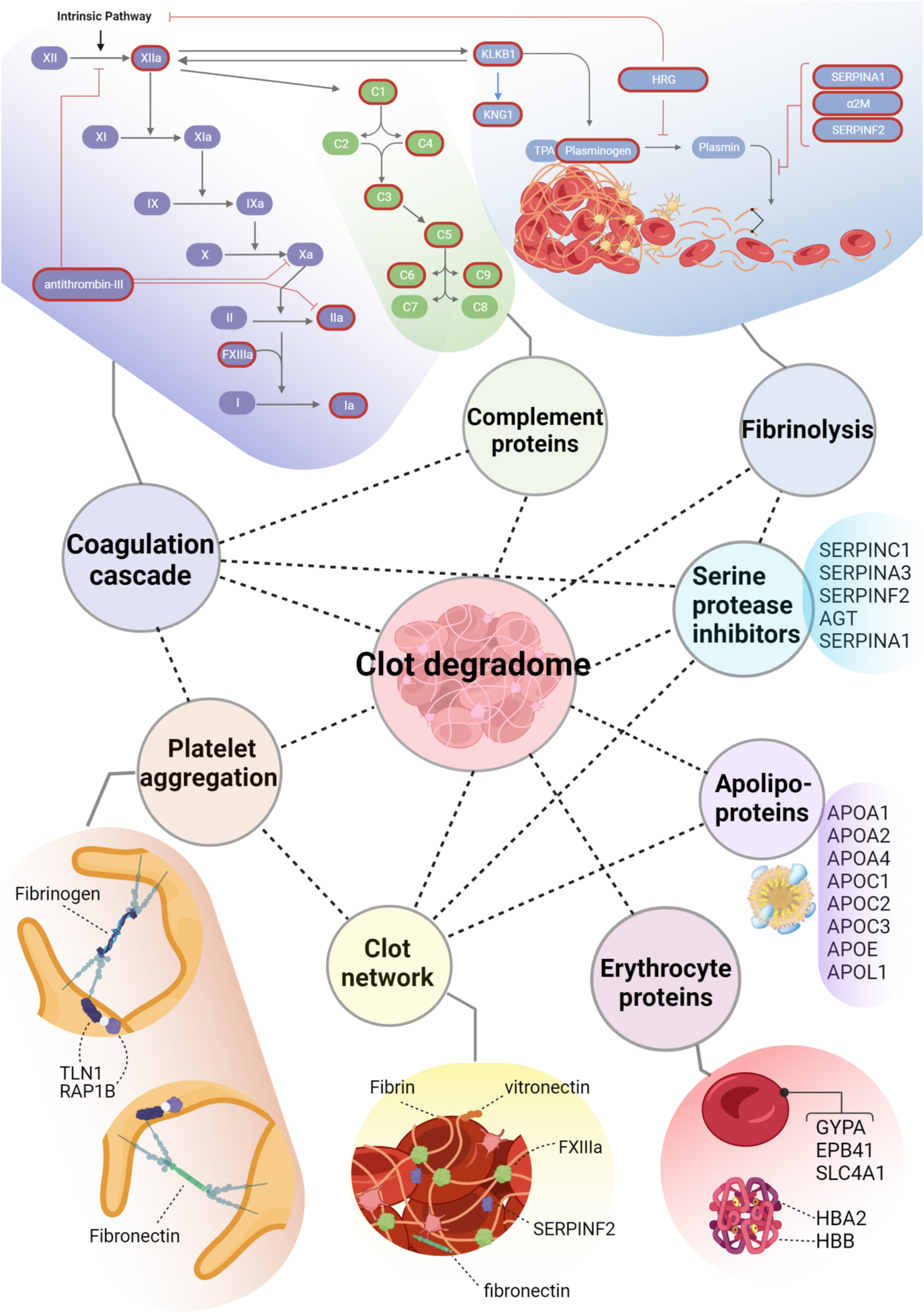
Snake venom clot degradome. Simplified gene ontology network of proteins identified in the snake venom blood clot degradome. Protein targets associated with the coagulation cascade, complement system, and fibrinolysis are circled in red, all other targets are listed in black.

Using Peptigram software, we characterized the unique protein cleavage patterns for all three fibrinogen subunits, noting enhanced proteolytic activity and broader coverage across all fibrinogen subunits for BIAR and CRAT compared to all other species. The cleavage of fibrinogen chains alpha, beta, and gamma with varying specificities by venom proteases has repeatedly been observed (16,54). We also noted varying cleavage patterns across alpha, beta, and gamma fibrin subunits, with the most thrombolytic venoms targeting regions essential for clot stability, including regions involved in cross-linking, fibrinolysis inhibition, and cell adhesion. Overall, it appears that the increased thrombolytic activity from BIAR and CRAT venoms is due to their enhanced ability to target multiple sites across all fibrin chains critical to clot stability and structure.

Venoms appear to target various cross-linking and/or binding sites across fibrin chains to elicit thrombolysis. Alpha chain cross-links across the aC domains are integral to fibrin polymer assembly (57). We found extensive degradation across alpha chain interchain binding sites, particularly by BIAR and CRAT, with a high number of peptides originating from the αC domain, indicating possible compromise of alpha cross-links and therefore clot stiffness and elasticity. BIAR and CRAT also produced multiple cleavage sites at or flanking plasmin cleavage sites in the alpha and beta chains. This plasmin-like activity is likely unrelated to endogenous plasmin activity, as we did not detect peptides associated with plasminogen activation. These thrombolytic venoms also uniquely targeted residues flanking the alpha-2-antiplasmin binding site of chain alpha, possibly neutralizing the protective effects against fibrinolysis of alpha-2-antiplasmin binding. BIAR and CRAT showed significantly higher degradation of fibrinogen chain beta also, particularly in the beta-chain polymerization site towards the N-terminal where binding occurs to the distal site of another fibrin strand.

The gamma chain of fibrin is pivotal to the formation of structurally sound fibrin polymers with high mechanical resistance, and fibrin polymers lacking gamma chain crosslinks produce unstable clots that easily fragment (58). Fibrinogen gamma also accelerates thrombin-mediated factor XIII activation, enhancing clot stability mediated by factor XIIIa activity. We specifically noted cleavage by BIAR and CRAT of the C-terminal portion of fibrin chain gamma (16), the region involved in fibrin stabilization via cross-links (59), and platelet interactions via integrin binding (60–64). As platelet aggregation is another critical step in clot formation, the targeting of this region by CRAT and BIAR venoms likely plays a significant role in thrombolysis by compromising the stiffness and structure of the fibrin network and by interfering with platelet aggregation in stable clots.

Coagulation factor XIIIa is critical for clot stabilization during coagulation via its catalysis of fibrin cross-links and of cross-linking of fibrin to other proteins including alpha-2-antiplasmin (65,66), inter-alpha-inhibitor (67), fibronectin, HRG (68), ECM1, a2M, plasminogen, and c3 (69). We identified cleavage of factor XIII A subunit in all venoms, which remains associated with the clot after the B subunit is released (70). We also identified the activation region in clots treated with all venoms but a higher abundance in BIAR-treated clots, as well as downstream degradation products of factor XIIIa from the clot degradomes of BIAR and CRAT. There is evidence that the activation peptide is not cleaved during activation of FXIII subunit A (71), and therefore its cleavage and further degradation by CRAT and BIAR may have affected FXIIIa-mediated clot stabilization.

Enzyme-rich thrombolytic venoms also demonstrated increased activity towards other clot-stabilizing proteins that are substrates of FXIIIa (66), including fibronectin, vitronectin, ECM-1, and lumican (16,19,20). Fibronectin covalently cross-links to fibrin to significantly increase clot stability and size (72,73) and association with fibronectin specifically has been shown to increase the adhesive properties of fibrin, making fibronectin critical for clot retention of stabilizing platelets and other cellular material (74,75). We noted cleavage sites flanking cell-attachment and Fibulin-1 (FBLN1) binding sites of fibronectin in the clot degradome of multiple species, but only BIAR and CRAT clot degradomes showed peptides from the N-terminal region of fibrin-binding site 2. Therefore, venom may compromise the stabilizing properties of fibronectin-mediated cell and platelet adhesion to clots by degrading both fibronectin sites of cell adhesion and interfering with fibrin-fibronectin interactions.

Vitronectin (76,77) is known to inhibit fibrinolysis by complexing with type 1 plasminogen activator inhibitor on fibrin polymers (74,78,79). Further, vitronectin preferentially binds to the carboxyl-terminal of the gamma chain of fibrin, another venom target discussed above involved in clot stabilization via cross-linking (77). Vitronectin-deficient thrombi are inherently unstable and show an increased tendency to embolize (77,80). BIAR and CRAT had the highest intensity of peptides resulting from vitronectin degradation, and we noted peptides that spanned the RGD cell attachment site of vitronectin in NASA, CRVI, and CRAT, suggesting further interference with structural protein-mediated cell adhesion.

The comprehensive understanding of how various plasma proteins contribute to thrombus formation, structure, and stability by altering the physical properties of thrombi is pivotal for identifying novel targets of future anticoagulant and thrombolytic therapies (81). The clot-based proteins discussed above that represent targets of venom proteases support clot-stabilization via structural changes that determine the physical properties of a thrombus including clot permeability, resistance to lysis, clot stiffness, and cell-adhesive properties (56,57,75). Measurable structural differences in clot architecture caused by deficiencies or enrichment in specific clot-bound proteins can be detected in proteomic clot profiles and correlated with various disease states and clinical outcomes (82–84), allowing for the identification of therapeutic targets (85).

Though snake venoms are toxic mixtures with a wide array of biological effects, several proteins are pharmacologically active but nontoxic when isolated (86,87). Numerous snake venom toxins have already been identified for their therapeutic utility in the treatment of coagulopathies (88–90), For example, Integrilin®, a peptide derived from *Sistrurus miliarius barbouri* venom, is used to treat acute coronary syndrome due to its effects on platelet aggregation (91,92). Further, venom proteins specifically have shown utility as diagnostic research tools in the study of hemostatic disorders (8). For example, venom enzymes that activate coagulation factors or other plasma proteins (93,94) have been used to test for specific protein levels in blood and to elucidate the mechanisms behind the coagulation cascade (8,95). Therefore, cataloging the entire suite of toxin targets in blood clots and the resulting functional effect of their cleavage has tremendous implications for the bioprospecting of thrombolytic therapies.

## Acknowledgments

We thank Benjamin German, Max Hicks, Eric Gren, Jason Folt, and Brian Wilson for their assistance in the field.

## Funding

This work was supported by Department of Defense (DoD) grant ID07200010-301-35 to A.J.S, C.F.S, and N.P.B. and the Fulbright U.S. Scholar Grant to C.F.S.

## Notes

### Competing Interest Statement

The authors have declared no competing interest.

