## Supplementary figures and images for "Discerning Specific Thrombolytic Activities and Blood Clot Degradomes of Diverse Snake Venoms with Untargeted Peptidomics"

### Supplemental Figure 1

This website uses cookies to ensure you get the best experience on our website Learn more
Got it!

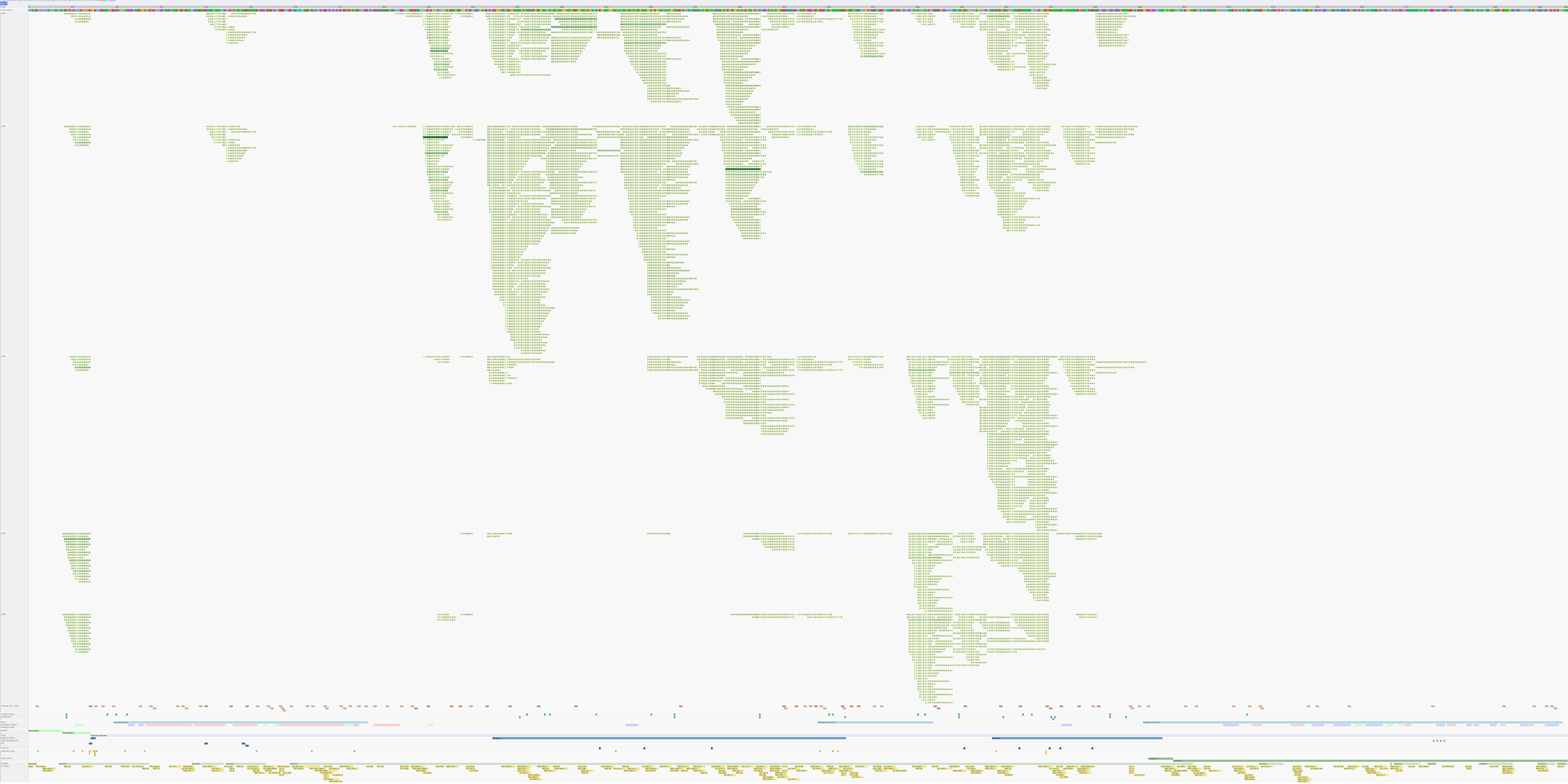

### Supplemental Figure 2

This website uses cookies to ensure you get the best experience on our website. Learn more

Got it!

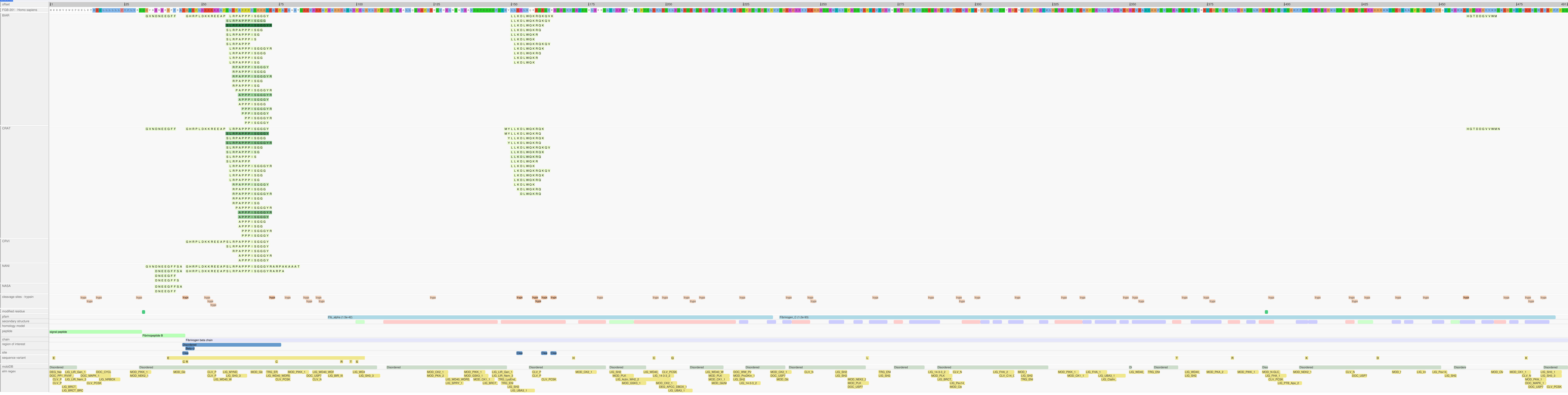

### Supplemental Figure 3

This website uses cookies to ensure you get the best experience on our website.Learn more
Got it

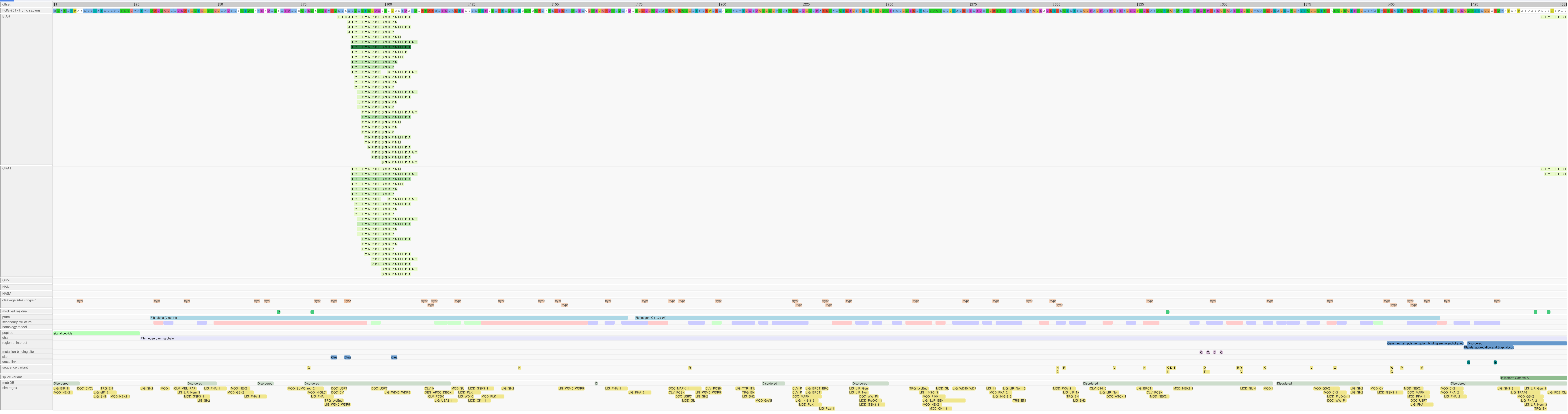
